# Decoherence via Demyelination Hypothesis (DDH): A Mechanism of Cognitive Decline During Aging

**DOI:** 10.64898/2026.05.22.727307

**Authors:** Iosif M. Gershteyn, Nikola T. Markov, Molly Olzinski, Joel Kramer, Kaitlin B. Casaletto, Lisa M. Ellerby, David Furman

**Affiliations:** Medical University of South Carolina, Charleston, SC, USA; Ajax Biomedical Foundation, Newton, MA, USA; ImmuVia Inc. Cambridge, MA, USA; Buck Institute for Research on Aging, Novato, CA, USA; University of California, San Francisco, Weill Institute for Neurosciences, Memory and Aging Center, Department of Neurology, San Francisco, CA, USA; Stanford 1000 Immunomes Project, Stanford University School of Medicine, Stanford, CA, USA

## Abstract

Progressive cognitive decline and loss of white matter integrity are observed during aging, but whether these two processes are connected remains unclear. We propose the Decoherence via Demyelination Hypothesis (DDH) as a mechanism linking non-uniform, tract-specific myelin loss during aging, which results in the degradation of conduction timing that is required to coordinate long-range neuronal assemblies. This impairs the dynamic assembly of task-dependent functional networks and promotes age-associated cognitive change. Axonal myelination determines conduction velocity as well as signal transmission fidelity, properties essential for phase-locked integration of long-distance inputs with local oscillations and for the assembly of mesoscale functional brain networks. If myelin loss is heterogeneous across tracts, the resulting timing perturbations should be heterogeneous as well, with disproportionate impact on networks supporting higher-order cognition. To test this, we analyzed structural and microstructural MRI in 638 individuals aged 40–99, quantifying fasciculation in different white matter pathways as well as neurite orientation dispersion and density imaging (NODDI) to the gray–white matter interface beneath cortical regions. We observed tract-specific, non-uniform myelin decline, with significant nonlinear losses in tracts serving high order cognition and memory (uncinate fasciculus, fornix, corpus callosum), consistent with accelerated late-life vulnerability in pathways implicated in cognitive aging and increased risk for neurodegenerative diseases. Tract-level degeneration tracked with age-related shifts in network organization and cognition. These findings may support DDH as a framework for age-associated cognitive change. Particularly, tract-specific timing perturbations drive the breakdown of coordinated network activity in aging.

## Introduction

Neural processing is based on the spatial and temporal patterning of the activity in distributed neuronal populations. However, early observations of focal lesions led to the view that cognitive functions are localized to specific cortical areas (Lanska, 2016). Anatomical evidence suggests that the structure of cortico-cortical connectivity is characterized by a dense network of inter areal connections, thus providing a substrate for coordinated parallel processing (Markov et al., 2013). This parallel processing is evidenced by neuronal populations from different brain areas entering synchronized activity (Nowak et al., 1999). High-density single-unit neuronal recordings during complex behaviors show that these behaviors are supported by a highly distributed neural activity spanning multiple brain regions (Zhang et al., 2025). Studies of fMRI activations during different cognitive tasks demonstrate that brain operations recruit coordinated activity across multiple large-scale networks, including the frontoparietal or multiple-demand network, dorsal and ventral attention networks, limbic and medial temporal lobe networks and sensorimotor circuits, where higher activation associates with better performance (Assem et al., 2020; Cai et al., 2024; Fox et al., 2005; Mione et al., 2026). Brain functions, including executive control, attention, working and episodic memory, language, and motor behavior, are now understood as emergent properties of interactions within and between widely distributed large-scale networks rather than being localized to single cortical centers (Buzsáki & Draguhn, 2004; Combrisson et al., 2025; Fries, 2005; Papadimitriou & Friederici, 2022; Wang, 2010).

Consistent with the timing function, the emergence of coherent large-scale network activity depends critically on the structural properties of the white matter pathways linking distant cortical and subcortical regions (Chu et al., 2014; Nelson et al., 2025; Salami et al., 2003; Vandewouw et al., 2021). Myelin sheaths increase signal precision by accelerating axonal conduction velocity and reducing temporal jitter in signal propagation, thereby aligning the arrival times of action potentials at target circuits and preserving the phase relationships necessary for inter-regional oscillatory synchrony (Fields, 2008; Kim et al., 2013; Pajevic et al., 2014). This is evidenced by the protracted parallel development of human cognition and myelination (Buyanova & Arsalidou, 2021; Corrigan et al., 2020; Konrad et al., 2013; Miller et al., 2012). The maturation of myelin during adolescence has been directly connected to the maturation of neurophysiological networks (Hunt et al., 2016; Kwon et al., 2020; Park et al., 2022). Alternatively, it has been observed that the three waves of myelin changes: establishment, maturation, and atrophy, coincide with the ages of onset of the neurodevelopmental, psychiatric, and neurodegenerative diseases (de Faria et al., 2021). The connection between myelin and neurological diseases is strengthened by the fact that genes associated with these conditions are highly enriched in oligodendrocytes (OL) (de Faria et al., 2021).

Mylenination patterns are not fully predetermined. A growing body of evidence suggests an interplay between cognitive activity and myelination. While oligodendrocytes have been demonstrated to build myelin sheaths around inert axon-shaped objects, they also generate “smart wiring” leading to active axons becoming more myelinated (Bechler et al., 2018). Early myelination involves repetitive ensheathment of axons that resolves through a low stabilization rate, supporting the idea of activity-dependent myelin plasticity (Almeida & Macklin, 2023; Fields, 2015; Pease-Raissi & Chan, 2021). Activity-dependent plasticity is likely a fundamental property of oligodendrocytes in stabilizing active neural circuits via a feedback loop of neuronal activity increasing oligodendrogenesis, maturation of oligodendrocytes, and densifying myelin sheaths (Bercury & Macklin, 2015; de Faria et al., 2018). The importance of oligodendrocytes in learning has been demonstrated by showing that both increased proliferation during learning of motor skills and the inability to learn new motor skills when differentiation of OLs is blocked (Aye et al., 2026; Birey et al., 2017). High-order cognitive functions such as working memory and episodic memory require generation of myelinating oligodendrocytes (Barboza et al., 2022; Munyeshyaka & Fields, 2022; Shimizu et al., 2023). Together, these findings establish myelination as a form of experience-dependent plasticity.

By adapting conduction velocities to experience, myelin plasticity provides an additional flexible system that complements the well-known Hebbian plasticity. Hebbian plasticity operates at the vertex level of neuronal networks while myelin plasticity operates on the edges of the network. In this respect, Hebbian plasticity establishes fine-grain communication weights controlling memory, while adaptive myelination facilitates the flexible establishment of dynamic function-specific neuronal assemblies. Adaptive myelination appears to be necessary to build the complex patterns of oscillations that establish cohesive functional networks in the brain (Flower et al., 2025; Mount & Monje, 2017; Pajevic et al., 2014, 2023). If this is true, then disruption of the myelination patterns in the brain should produce cognitive consequences. Here, we propose the Decoherence via Demyelination Hypothesis (DDH) and provide structural evidence consistent with its core prediction: that heterogeneous, tract-specific myelin loss during aging selectively disrupts the conduction timing of long-range projections, degrading the inter-regional neuron communication coherence necessary for task-dependent network formation and thus produces the cognitive deficits characteristic of aging.

## Results

### Decoherence via Demyelination Hypothesis

DDH proposes a specific mechanistic chain: heterogeneous age-related demyelination of long-range white matter projections disrupts conduction timing, degrades inter-regional neuronal communication coherence, and produces the cognitive deficits characteristic of aging. The physical embedding of the brain network and the non-instantaneous travel times of signals along axons constitute a constraint to long-distance inter-neuronal communication. All things being equal, two simultaneous signals sent at the same time from a distant and a closer brain area will not reach their target area at the same time, a condition required for their integration. A well-calibrated increase in myelination of the longer pathway will speed up the signal along its tract and ensure that both the signal traveling a longer distance and the one traveling a shorter one reach their target at the same time (**Figure 1A**). This tuning emerges from a feedback loop in which oligodendrocyte precursor cells (OPCs) respond to neural activity by differentiating into oligodendrocytes, adapting white matter structure to cognitive demand (**Figure 1B**). During aging under heterogeneous demyelination of long-range tracts, coherent signal arrival degrades for signals travelling from different locations (**Figure 1C**). Testing the DDH hypothesis structurally requires establishing three observations: i) that white matter integrity declines with age in a tract-specific, heterogeneous manner, preferentially affecting higher-order association pathways; ii) that this decline reflects changes in the cellular compartments consistent with myelin loss rather than axonal loss; and iii) that these microstructural changes covary with cognitive performance in a pattern consistent with distributed network dysfunction. We address each prediction in turn using diffusion-weighted MRI in 638 adults aged 40–99 years from the UCSF Brain Aging Networks for Cognitive Health (BrANCH) cohort.

**Figure 1.**
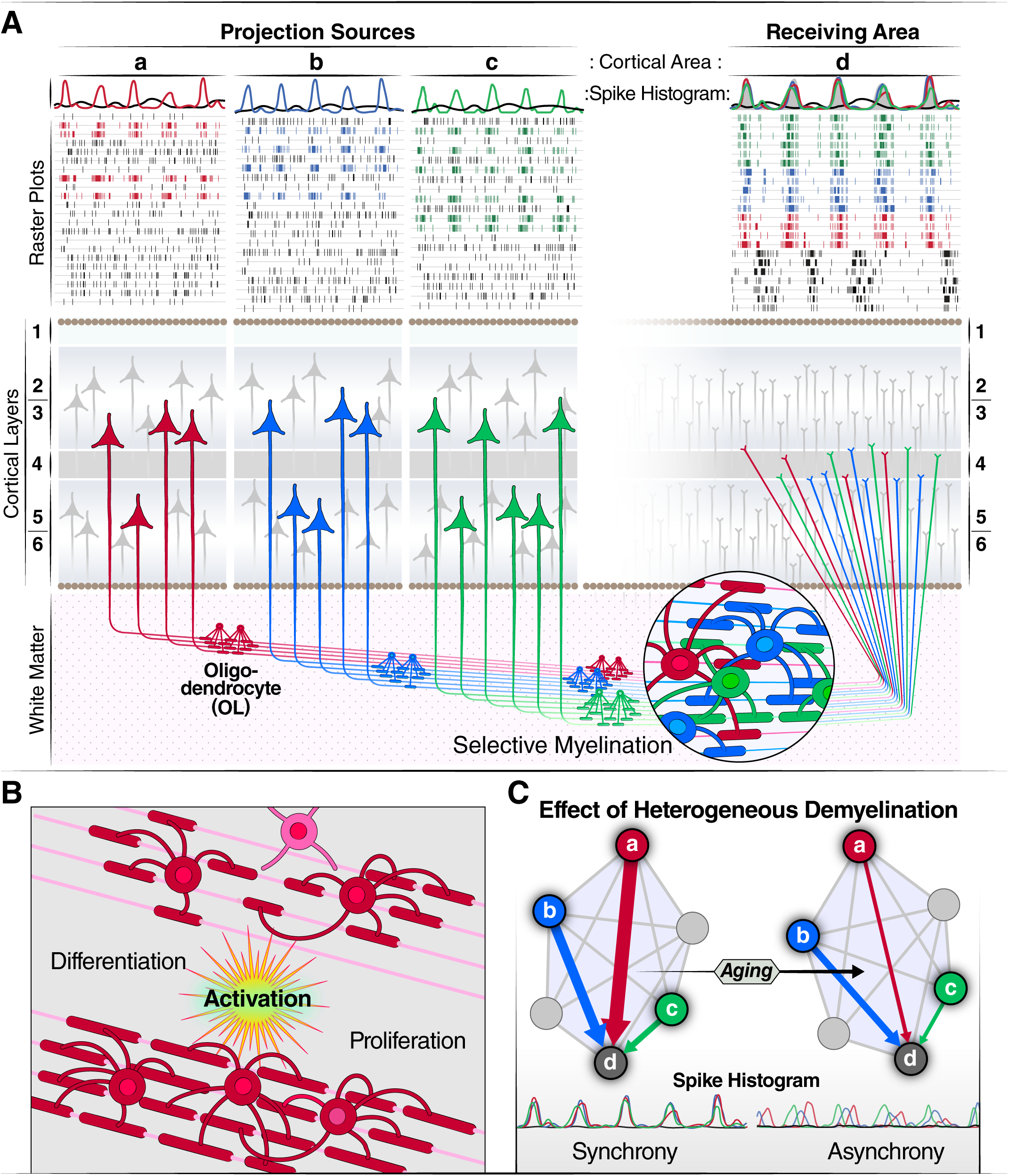
The Decoherence via Demyelination Hypothesis (DDH) — a conceptual model. **(A)** Neuronal ensembles from different brain areas communicate through coherence. Myelin patterns in the white matter compensate for the difference in distance between brain areas by controlling conduction velocities. Simultaneous activations arising in phase with synchronized local field potentials arrive simultaneously in a target area. Selective myelination allows for tuned conduction velocities. **(B)** Cellular basis of conduction velocity tuning: oligodendrocyte precursor cells (OPCs) respond to neural activity by differentiating into myelinating oligodendrocytes (OLs), creating a feedback loop between cognitive demand and white matter structural adaptation. **(C)** DDH predicts that heterogeneous demyelination of long-range white matter tracts disrupts the timing of inputs to target circuits. Coherent, synchronized signal arrival (left) degrades to temporally dispersed, incoherent inputs (right), impairing the formation and persistence of task-dependent functional networks. Higher-order association regions (e.g. HPC, PFC, ATL, IPS, ACC) connected by late-myelinating tracts are predicted to be the most vulnerable. HPC, hippocampus; PFC, prefrontal cortex; IPS, intraparietal sulcus; ATL, anterior temporal lobe; ACC, anterior cingulate cortex.

First, we measured the mean fractional anisotropy (FA) and mean diffusivity (MD) in a set of white matter tracts, and we explored the dynamic changes of fasciculation of white matter pathways associated with aging (Madden et al., 2009). Second, we interrogated the microstructure of the superficial white matter, targeting the highly complex transitional zone immediately beneath the cortical mantle described in recent high-resolution mapping studies (Hwang et al., 2026). We applied a NODDI model with evaluation of the different intra and extracellular compartments and neurite density and orientation. (Sone, 2019; Song et al., 2025). Third, we use canonical correlation analysis (CCA) to identify which compartment changes over age, are associated with changes in cognitive functions. Our results are described below.

### White matter integrity declines heterogeneously with age, with accelerated loss in higher-order association tracts

To characterize the relationship between age and white matter integrity in adults aged between 40 and 99 years, we first examined the magnitude and direction of age effects on FA and MD, across 28 major white matter tracts in 638 participants using robust linear models adjusted for sex (**Supplementary Figure 1**). Both FA and MD demonstrated significant age-related changes across multiple white matter pathways. FA reflects fasciculation i.e. the directional orientation and alignment of axonal fibers. For FA, the strongest negative age effects were observed in limbic pathways, including the column and body of fornix (β = -0.0026), which is critical for episodic memory formation and spatial navigation. Association fibers connecting frontal and temporal regions showed similarly pronounced effects, including the uncinate fasciculus (β = -0.0027), its role spans social-emotional processing, memory, language, motivation and reward-based learning. The anterior limb of internal capsule (β = -0.0024), a projection pathway carrying thalamo-cortical fibers and involved in executive control and decision making, also demonstrated substantial age-related FA decline. Commissural fibers, particularly the body of corpus callosum and genu of corpus callosum, showed moderate to strong age effects, reflecting changes in interhemispheric communication. Primary motor projection pathways such as the corticospinal tract and cerebellar peduncles exhibited relatively smaller effects.

MD measures the compaction of the tracts, composed of axonal fibers and the sheaths of compacted myelin that surround them. Denser oligodendrocytes are reflected in lower MD values, while expansion of the extracellular spaces would lead to increased values. MD showed positive age effects across all tracts, with the most pronounced increases in association fibers, such as the uncinate fasciculus (β = 0.010). Limbic pathways showed similarly strong effects, including the fornix (β = 0.008). Commissural fibers of the corpus callosum also demonstrated substantial MD increases (genu: β = 0.007; body: β = 0.006; splenium: β = 0.007). Thalamo-cortical projection pathways showed moderate increases, including the anterior limb of the internal capsule (β = 0.006), and anterior corona radiata (β = 0.008). Primary motor projection pathways, including the corticospinal tract and middle cerebellar peduncle, demonstrated the most modest age-related increases in MD. Taken together, these results indicate a dramatic age-dependent reorganization of the white matter tracts reflected in both their compaction and the homogeneity of the dominant orientation of the fiber pathways.

We next examined whether age-related changes in FA followed linear or nonlinear trajectories across the the age range (**Figure 2**). We built linear models and models with polynomial terms that would reflect the acceleration of the age effect over time. Model comparison using Akaike Information Criterion (AIC), supported by a weight of evidence for curvature estimate (ω), revealed substantial heterogeneity in trajectory shapes across white matter tracts. Fifteen tracts were best fit by nonlinear models (ω nonlinear > 0.77), with the strongest support observed in association fibers and limbic pathways: anterior limb of the internal capsule (ΔAIC = -26.88, ω nonlinear = 1.00), uncinate fasciculus (ΔAIC = -26.65, ω nonlinear = 1.00), and splenium of the corpus callosum (ΔAIC = -26.08, ω nonlinear = 1.00). These nonlinear trajectories indicate relative stability or gradual decline in early to middle adulthood, followed by accelerated decline after approximately age 60, as exemplified by the hill and valley pattern in the uncinate fasciculus. In contrast, eight tracts best fit linear decline models, including the anterior corona radiata and genu of the corpus callosum, suggesting more uniform degradation across the adult lifespan. Notably, five tracts showed no significant age effects, including the internal capsule posterior limb, which exhibited stable FA values across all age groups. MD exhibited more uniform age-related patterns across white matter compared to FA (**Supplementary Figure 2**). All 28 examined tracts demonstrated significant nonlinear relationships with age (all ω nonlinear ≥ 0.93), characterized by gradual increases in early adulthood that accelerated after approximately age 60. The strongest evidence for nonlinear trajectories was observed in association and interhemispheric fibers, including the Internal capsule anterior limb (ΔAIC = -64.83, ω nonlinear = 1.00), Uncinate fasciculus (ΔAIC = -29.07, ω nonlinear = 1.00), Splenium of corpus callosum (ΔAIC = -42.53, ω nonlinear = 1.00). To confirm the model fit is consistent with the local trends in the data we plotted the average values of MD and FA in 5-year bins. Visual inspection confirmed consistent j or u-shaped patterns. This universal nonlinear pattern suggests that age-related increases in water diffusivity, potentially reflecting myelin degradation and increased extracellular space, follow a consistent temporal progression across diverse white matter systems regardless of their functional specialization.

**Figure 2.**
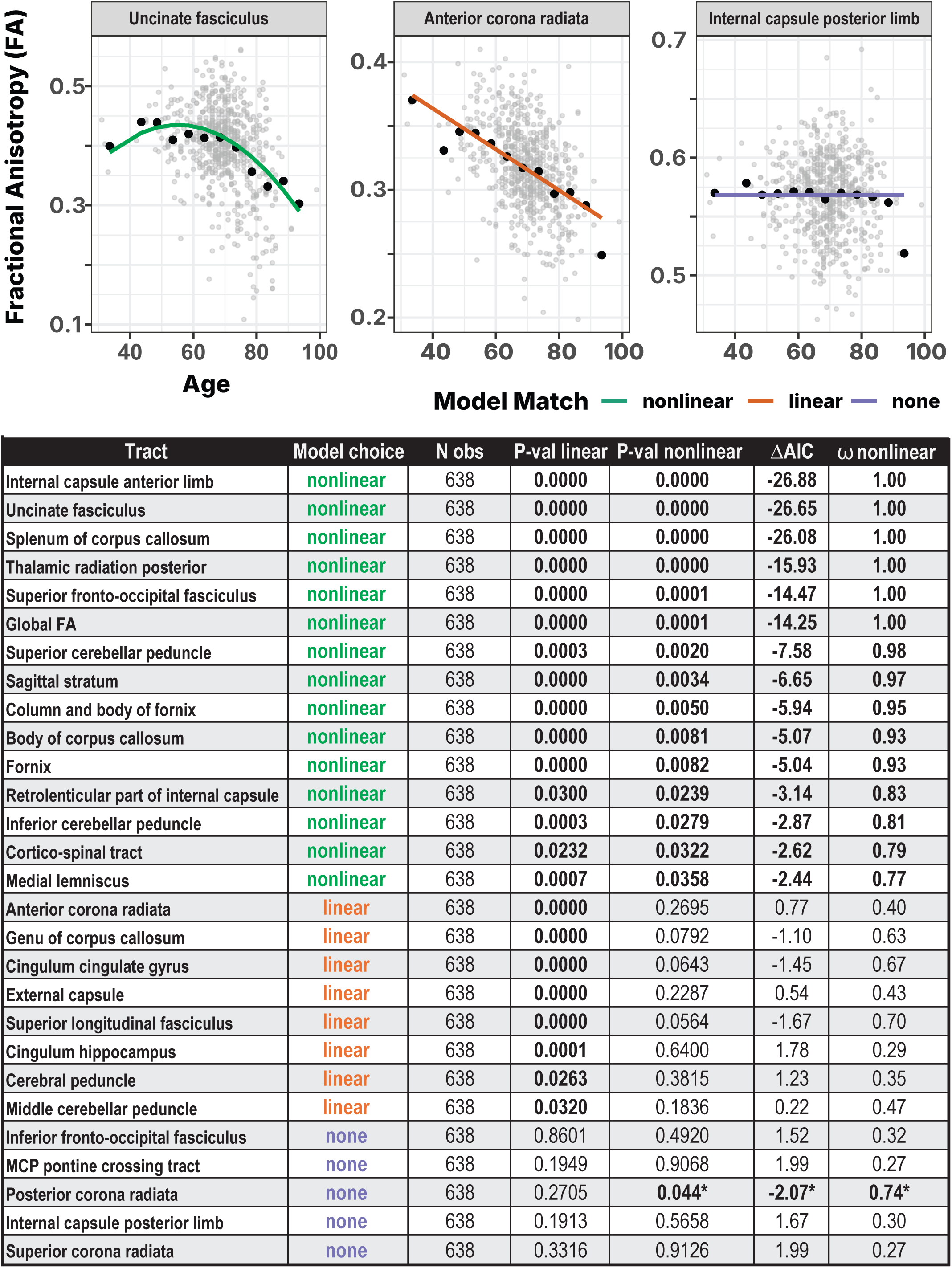
Age-related trajectories of fractional anisotropy across major white matter pathways show heterogeneous accelerations. **(A)** Representative scatter plots of mean tract values from first visit dwMRI scans (gray points) FA against chronological age. Large black points indicate mean FA within 5 years interval. For each tract, age trajectories were estimated using sex adjusted regression models; the best fitting model was selected using a combined statistical and information-theoretical criterion (see Table in panel B). Colored curves denote the chosen model for each tract: green = nonlinear (quadratic) age dependence; orange = linear age dependence; purple = non-detectable age effect. Panels are scaled independently to emphasize tract-specific dynamic ranges. **(B)** Table of model fitting diagnostics for each pathway. **Tract**: white matter pathway of interest. **Model choice**: best supported age model (nonlinear (green), linear (orange), none (purple), based on significance testing and information criterion evidence. **N obs:** number of individuals contributing FA measurements to that tract. **P-val linear**: significance score of nested ANOVA F test of the linear age effect (*FA∼Age + sex*) versus a gender-only null model (*FA ∼ sex*). **P-val nonlinear**: significance of nested ANOVA F test for curvature (quadratic term) (*FA ∼poly(Age, 2) + sex*) beyond the linear model (*FA∼Age + sex*); **ΔAIC**: Akaike Information Criterion (AIC) difference between linear and nonlinear term, if (ΔAIC ≤ –2, and/or ω ≥ 0.70) are labeled nonlinear; those with significant but non-curved age effects are labeled linear; and tracts without significant age dependence are labeled none. **ω nonlinear** Akaike weight for the nonlinear model, representing the relative probability that curvature provides the best description of age-related change.*Due to outliers leverage *Posterior corona radiata* reaches significance on the nonlinear age relationship but the linear model is non-significant, therefore we do not conclude on relationship with age.

### Age-related microstructural changes preferentially associate with extracellular water fraction, consistent with myelin rather than axonal loss

Measuring FA and MD provides an overall impression of the integrity of fiber tracts in the brain, but doesn’t capture projection specificity (multiple pathways converge in the same white matter tracts) and cannot disentangle the potential loss of axonal fibers from changes in the health of myelin support to these fibers. To obtain a more detailed determination of microstructural changes, we focused on NODDI (neurite orientation dispersion and density imaging) models of diffusion weighted imaging of the white matter cushions under 34 cortical brain regions. Robust linear modeling corrected for sex revealed distinct age-related patterns across the three NODDI parameters in white matter cushions underlying these cortical regions (**Figure 3**). Age demonstrated widespread and significant effects on both intracellular volume fraction (ficv) and free water fraction (fiso), while the orientation dispersion index (odi) showed minimal dependence on age. For ficv, age was negatively associated with intracellular volume across nearly all examined regions, with the strongest effects observed in temporal regions, including the banks of the superior temporal sulcus, inferior temporal cortex, and temporal pole (significant β ranging from approximately -0.002 to -0.001). Additional regions showing robust ficv decline included the entorhinal cortex, insula, anterior cingulate, and, interestingly subdivisions of the inferior frontal gyrus including BA44 and BA45 regions. Confidence intervals for most of these regions excluded zero, indicating statistically reliable effects. These negative associations indicate age-related reductions in the volume of neurites within white matter. Conversely, fiso exhibited predominantly positive associations with age across most brain regions, with effect sizes roughly mirroring the ficv findings in magnitude but opposite in direction (significant β ranging from 0.001 to approximately 0.002). The strongest age-related increases in free water were observed in the inferior frontal gyrus (BA44 and BA45) both involved in language production semantic processing, working memory and executive cognitive control, lingual involved in word recognition visual processing and memory, pericalcarine roles in visual processing, and the white matter underneath the cingulate gyrus (both anterior and posterior) involved in decision making, performance monitoring and attention regulation. This pattern suggests age-related accumulation of freely diffusing water, potentially reflecting increased extracellular space or reduced tissue density that could include loss of myelinating cells and myelin sheets.

**Figure 3.**
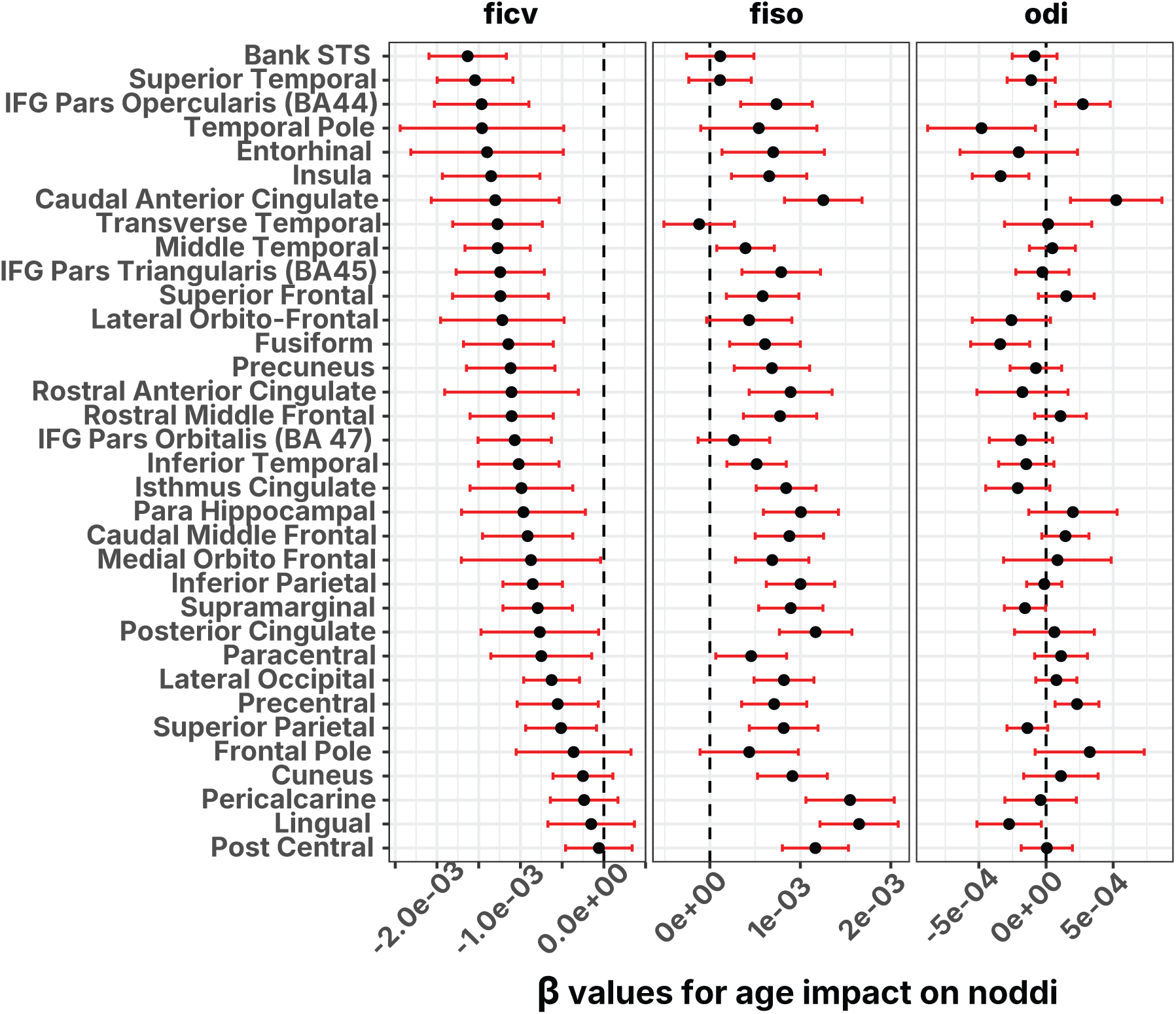
Population level effect of age on 3 NODDI models applied to the white matter cushions of thirty-four brain regions demonstrates that age impacts freely diffusing water (fiso) and intracellular volume (ficv) but not the neurite orientation dispersion index (odi). Brain parcels (y-axis) correspond to voxels of the white matter cushions under the corresponding gray matter region. Age effect coefficients (β age ± 95% CI) were estimated using robust linear models (NODDI value ∼ Age + sex). Points indicate β-estimates and red bars show confidence intervals and the dashed line marks zero effect. Parcels are ordered by the strength of the age effect in ficv, with fiso and odi shown in parallel facets.

In contrast to ficv and fiso, the orientation dispersion index (odi) showed substantially smaller and more variable effects of age. Beta estimates were tightly clustered near zero with confidence intervals frequently crossing zero, indicating minimal age effects on neurite orientation dispersion across the examined white matter regions. Visualizing the beta values for the three measures in parallel (ranked by ficv effect size) revealed spatial heterogeneity in age sensitivity, with some regions showing atrophy exclusively in ficv while others present significant changes both in ficv and fiso. Taken together the observations from cohort based diffusion weighted imaging showed that, i) there is a significant heterogeneous remodeling of white matter fiber pathways with age that accelerates especially after age 60 and is most drastic in pathways involved in higher order cognition with more moderate effects on sensory and motor pathways; ii) the changes impact less the fibers orientation and dispersion but are more related to modification of extracellular space and neurite packing quality that could be attributed to decrease of the volume occupied by supporting glial cells.

### White matter microstructure and cognitive performance share a single dominant age-dependent dimension driven by extracellular water accumulation

To investigate the multivariate relationship between white matter microstructure and cognitive function, we performed canonical correlation analysis (CCA) between NODDI-derived imaging metrics (ficv, fiso, and odi) across 34 white matter parcels and seven cognitive assessments matched within a one-year window of the imaging session. After excluding outliers and restricting to the first visit per participant, the analysis included 638 complete observations spanning ages 40-100 years. The CCA revealed a strong and highly significant association between brain structure and cognitive performance. The first canonical variate pair (CV1) explained 52% of the shared variance between imaging and cognitive domains (R = 0.72, p < 2.2×10⁻¹⁶; **Figure 4A, Supplementary Figure 3**). This dominant mode captured a systematic relationship wherein participants with greater white matter microstructural damage (lower U1 weights) exhibited poorer cognitive performance (lower V1 weights). Subsequent canonical variate pairs showed progressively weaker correlations (CV2: R² = 0.40; CV3: R² = 0.38), with only the first pair reaching statistical significance after FDR correction (**Supplementary Figure 3**). The strong age-dependence of this brain-cognition relationship was evident in the color gradient across the CV1 scatter plot (**Figure 4A**), with younger participants systematically distributed toward the upper-right quadrant (lesser microstructural damage and better cognition). Regression analyses confirmed that age significantly correlated with both canonical variates (U1: β = 0.041 ± 0.002, p < 0.001; V1: β = -0.058 ± 0.003, p < 0.001; **Supplementary Figure 3**), accounting for the majority of variance in each dimension. This pattern demonstrates that chronological aging represents the primary axis along which white matter microstructure and cognitive performance covary in this cohort.

**Figure 4.**
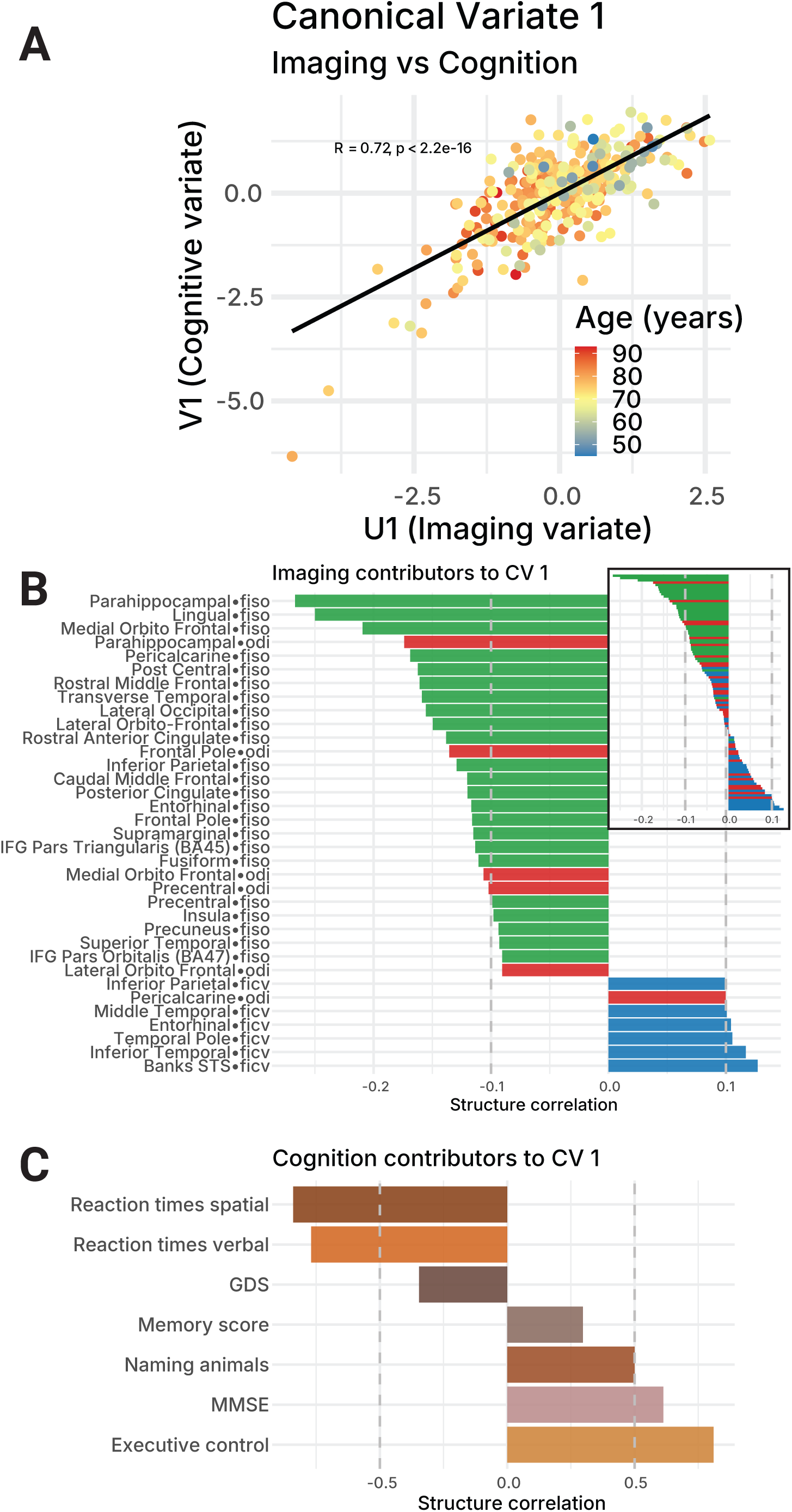
Age is a canonical variable between structural measures and cognitive evaluations. **(A)** Scatter plot of the canonical weights of the dominant pair of canonical variates 1 (CV1) colored by participant’s age. V1 is the cognition variate and U1 is the imaging variate. Patients with greater microstructural damage and poorer cognitive performance tend to be of more advanced chronological age. **(B)** Structural loadings of imaging features contributing to CV1. Upper right graph shows the full distribution of fiso (green), ficv (blue) and odi (red). The blow up represents the 35 top contributors to the CV1. Note that top contributors are mainly composed of fiso measures and represent higher cognitive regions. **(C)** Structural loadings of cognitive measures contributing to CV1. Top contributors include verbal and spatial reaction times as well as executive control score, performance on the Mini-Mental State Examination (MMSE) test exam and correct animal naming. Geriatric Depression Scale (GDS) and the Memory z-score are minor contributors.

Examination of the structural loadings revealed that CV1 was predominantly driven by isotropic water fraction (fiso) measurements from the cushions under the cortical regions (**Figure 4B**). Among the top 35 contributors to the imaging variate U1, fiso metrics constituted the majority of top imaging contributors, while orientation dispersion index (odi) showed more limited involvement, primarily through parahippocampal (odi: -0.16) and frontal pole regions (odi: -0.14). Regions with the highest contributing fiso parameters included the parahippocampal (fiso: -0.26), lingual (fiso: -0.25), and medial orbitofrontal (fiso:-0.21).

On the cognitive side, CV1 reflected a broad decline across multiple domains, with processing speed showing the strongest influence (**Figure 4C**). Reaction times for both spatial processing speed (loading: - 0.61) and verbal processing speed (loading: -0.55) tasks were the primary contributors, indicating that slowed information processing represented the most salient cognitive feature linked to white matter changes. Executive function (bsexzscore: 0.46) and global cognition as measured by MMSE (0.38) also loaded substantially on V1. Semantic retrieval performance (an_corr: 0.35) and episodic memory (memoryzscore: 0.26) showed moderate contributions, while depressive symptoms (GDS) exhibited only modest loading (-0.32), suggesting that mood disturbance was less central to the brain-cognition relationship captured by CV1. The multicollinearity among imaging metrics and among cognitive measures (**Supplementary Figure 3**) suggests that CCA successfully identified a coherent aging signature spanning both domains. The imaging variate captured a distributed pattern of white matter vulnerability affecting association cortex connectivity, while the cognitive variate reflected a general factor of cognitive aging weighted toward executive control and processing speed, consistent with distributed frontal connectivity dysfunction in aging. Taken together, these findings provide structural evidence consistent with the three core predictions of DDH: tract-specific heterogeneous white matter decline preferentially affecting higher-order association pathways, microstructural changes consistent with myelin compartment loss, and a dominant brain-cognition axis driven by extracellular water accumulation in regions subserving executive and memory functions

## Discussion

We sought evidence for the Decoherence via Demyelination Hypothesis, which posits that age-related cognitive decline arises from progressive demyelination of axonal tracts, disrupting the coherent network activity required for cognition. We observed heterogeneous breakdown of fasciculation across major white matter tracts measured by FA and accelerating age-related increase of mean diffusivity, consistent with a reduction of the white matter. These changes on the macro level of the interareal network are supported by analysis of the superficial white matter directly under brain regions where the long-distance axonal projections from the gray matter enter the white matter before joining the major fiber pathways. In these brain regions, NODDI models indicated a significant age relationship between the extracellular water compartment and the intracellular water. There was a modest age effect on the neurite orientation model, supporting the idea that the main effect of aging is an impact on the myelin integrity rather than loss of axons. Multivariate analysis further identified aging as the dominant latent variable linking superficial white matter decline to deficits in higher-order cognition.

Previous studies analyzing large datasets from the UK biobank, using linear models, have identified a heterogeneous alteration of FA and MD associated with aging (Isaac Tseng et al., 2021). Similar to our observation of brain-wide acceleration of MD increase after age 60, free water fraction in white matter is known to increase non linearly (Pieciak et al., 2023). Traumatic brain injury meta-analysis demonstrated specific cognitive outcomes related to attention and working memory associated with lower FA and higher MD in white matter tracts such as the corpus callosum, fornix, uncinate fasciculus, and the internal capsule (Wallace et al., 2018). Longitudinal follow-up of 123 participants show that lower myelin content at first scan is associated with faster cognitive decline (Gong et al., 2023). Combined with these studies, our results highlight implications for the way structural connectivity evolves with aging and the potential impact it can have on functional networks (**Figure 1C**). While the age-related changes of the white matter are quite drastic, the cognitive deficits associated with aging are mild in most people, indicative of compensatory mechanisms preserving brain dynamics against the structural decline (Chakraborty et al., 2025). Our results point to an accelerated demyelination in specific pathways, which would lead to asymmetry in the age-related changes of functional sub-networks thus hampering the ability of the compensatory mechanisms to “re-equilibrate” the aging brain networks.

Because multiple interareal projections share the same fiber tracts before fanning out to their respective targets, tract-level analyses conflate the effects on many edges of the functional networks. We sought to achieve more specificity to cortical regions by analyzing the gray-white matter interface under 34 cortical regions. We used neurite orientation dispersion and density NODDI model, which provides a more detailed insight on the microstructural properties of the regions of interest, fiso tracks the extracellular free water, ficv measures the intracellular water and reflects the compaction of the neurites while odi is an index of neurite orientation dispersion. NODDI measures have previously been identified to reflect aging and to correlate with cognitive performance (Beck et al., 2021; Kamiya et al., 2020; Raghavan et al., 2021). Most of the current studies have explored their relevance as clinical markers in measures of gray matter (Yu et al., 2024). Recently, NODDI has been linked to measures of cognitive impairment (Sathe et al., 2026). Our data confirmed that fiso and ficv are highly sensitive to aging for the white matter under most nodes of the cortico-cortical network while odi is only marginally impacted by normal aging. From the perspective of distributed functional networks, this result provides insight into the vulnerabilities of white matter pathways’ integrity to the aging process.

Our next step explored how the various changes in NODDI measures associate with cognition (Venkatesh et al., 2020). Different cognitive domains are impacted differentially by age, and taken together as a network view, there is a tendency toward dedifferentiation of psychological and cognitive dimensions (Vicentin et al., 2026). We performed a canonical correlation analysis against measures of cognitive performance and found that the major axis of correlation between cognition and brain structure is explained by age. Executive control and reaction time had the highest loadings in the first canonical variate, reciprocated by fiso and ficv loadings on the structural side.

There are some limitations in our study. In particular, we focused on the age-related atrophy of white matter pathways. While we describe the mechanisms that impair conduction along the axonal pathways linking distributed cortical regions, it is important to acknowledge that aging leads to significant changes within the cortical gray matter itself. The gray matter constitutes the nodes of the functional networks, whose coherence we propose is being undermined. Recent studies have explored the microstructural changes in cortical gray matter of different functional areas across the lifespan and found correlations with cognitive change that are similar to those found in our study of white matter (Merenstein et al., 2026). Changes in other parameters also indicate active remodeling in the gray matter (Acosta-Franco et al., 2025; Markov et al., 2022). Crucially, the atrophy of gray matter in normal aging is not primarily a consequence of neuronal death. Stereological studies have established that neuron counts in neocortex and hippocampus remain largely stable across the lifespan, with neuronal loss being a hallmark of neurodegeneration rather than normal aging (Morrison & Hof, 1997). Instead, the cortical decline appears to be the progressive loss of synaptic connections (Dickstein et al., 2013). Our data do not allow us to disentangle the directionality of these parallel processes at the level of individual participants. However, on theoretical grounds, the two forms of decline are unlikely to be independent: the degradation of long-distance myelinated pathways that we document here would be expected to reduce and desynchronize the activity arriving at cortical targets, progressively depriving local circuits of the coherent input that sustains synaptic maintenance and plasticity. White matter demyelination may be an upstream driver that accelerates synaptic pruning in downstream cortical regions. Future work involving functional and structural measures will be necessary to evaluate this hypothesis directly.

The Decoherence via Demyelination Hypothesis rests on two mechanistic pillars. The first is that cognition emerges from the dynamic assembly of distributed neuronal ensembles whose composition is reorganized in a task-dependent manner, a principle now well established in cross-areal recordings (MacDowell et al., 2025; Olson, 2026; Tafazoli et al., 2025), in the expansion and refinement of representational dimensionality preceding action selection (Kikumoto et al., 2024), in the rearrangement of network states underlying cognitive flexibility (Pagan et al., 2024), and in the reverberant large-scale dynamics that support distributed computation (Mejías & Wang, 2022). Such assemblies depend on precise inter-areal timing: hippocampal–prefrontal firing, for example, realigns to theta troughs during learning (Benchenane et al., 2010). The second pillar extends Fries’s communication-through-coherence framework to the multi-area case (Fries, 2005, 2015). Selective long-range communication requires that signals arrive in phase with local oscillations in the target area; when multiple projections must converge, this further requires that signals from sources at different distances arrive contemporaneously (**Figure 1A**). Conduction velocities must therefore be finely tuned across pathways, a tuning achieved by activity-dependent adaptive myelination shaped by repeated co-activation and learning (Bechler et al., 2018; Pease-Raissi & Chan, 2021). Together, these pillars predict that demyelination of long-range pathways during aging should misalign signal arrival times within functional assemblies, degrading their coherent activation and indeed, state switching and dynamic functional connectivity are impaired in aging and neurodegenerative disease (Arbabyazd et al., 2023; Salehi et al., 2019). DDH thus links a cellular process (heterogeneous myelin loss) to a network-level failure (loss of coherent assembly formation) and, ultimately, to the cognitive decline that defines aging.

## Materials and Methods

### Participants

The data for this study were derived from the UCSF Brain Aging Networks for Cognitive Health (BrANCH) cohort, established by the Memory and Aging Center at the University of California, San Francisco (UCSF). Participants for this cohort are primarily recruited via community outreach events, flyers, and media advertisements across the Bay Area in California. The research protocols have received IRB approval by the UCSF Committee on Human Research, and the researchers have followed the principles established by the Declaration of Helsinki. Written informed consent was obtained from all included participants. The data collected consisted of comprehensive health examinations, cognitive tests, and structural MRI scans. Scripts and data have been deposited at (Raw data repository ADKnowledge db synapse.org https://www.synapse.org/Synapse:syn75115498. Github scripts are found at https://github.com/NikolaTMarkov/DDH.

### Data Collection and Statistical Analysis

All statistical analyses were performed with R (version 4.4.1) (R Core Team, 2020). Deidentified data, scripts used for analyses, and other relevant documentation will be made available upon reasonable request by qualified researchers interested in replicating our results or performing independent analyses. Such requests should be sent to the corresponding authors.

### MRI Data Acquisition and Preprocessing

All brain MRIs were performed at the UCSF Neuroscience Imaging Center using either a Siemens Trio 3T or Siemens Prisma 3T scanner. Magnetization-prepared rapid gradient-echo (MPRAGE) sequences were used to obtain whole-brain T1-weighted images (Trio: TR/TE/TI = 2300/2.98/900 ms, flip angle = 9°; Prisma: TR/TE/TI = 2300/2.9/900 ms, flip angle = 9°). The field of view was 240 × 256 mm, with 1 × 1 mm in-plane resolution, 1-mm slice thickness, and sagittal orientation. Whole-brain diffusion MRI was acquired on Siemens platforms using multi-shell sampling suitable for diffusion tensor imaging (DTI) and neurite orientation dispersion and density imaging (NODDI). Standard preprocessing included correction for eddy currents, motion, susceptibility, and B0 field distortions.

### Tensor Model

FA and MD were computed using standard tensor fitting. Scanner- and protocol-related variation was harmonized using empirical Bayes ComBat across all sessions.

### NODDI Model

NODDI parametric maps were derived using established fitting procedures. Three metrics were retained: intracellular volume fraction (fICV), isotropic free water fraction (fISO), and orientation dispersion index (ODI).

### White Matter Parcellation

White matter regions-of-interest (ROIs) were extracted from atlas masks. Metrics were computed separately for left and right hemispheres. Preliminary analysis revealed that between-subject variability substantially exceeded within-subject hemispheric differences; therefore, left and right hemisphere values were averaged to produce a single tract estimate per participant.

### Neuropsychological Assessment

Cognitive assessments included bedside screening and information-processing tasks. Specifically: Mini-Mental State Examination (MMSE), Geriatric Depression Scale (GDS), memory z-score, animal naming test (correct responses), executive/bedside z-score, and processing speed (verbal and spatial reaction times).

### Data preprocessing

The overall database represents a comprehensive multi-visit assessment of physiological, structural and cognitive parameters with over 4K entries and 3K features. We developed a script that extracts all visits containing either cognitive assessments or MRI scans. Most MRI scans have a matching cognitive assessment on the same visit, but in the cases when no cognitive evaluation was made coincidently with the scan, we retrieved the nearest in time evaluation with a maximum window of 2 years from the scan date. When multiple evaluations satisfied this criterion, the temporally closest was selected. This rule avoided discarding potentially informative cognitive measurements while preventing physiologically implausible time intervals. Cognitive assessment sessions missing >35% of cognitive variables were excluded. Remaining gaps were imputed using median values from individuals of similar chronological age (within ±5 years), provided that at least three observations existed in the age neighborhood. This scheme preserved realistic age-dependent values and prevented distortion of multivariate structure. Outliers were screened consistently across DTI and NODDI. For patients with multiple MRI scan visits only one visit per patient was retained for the analyses. For tracts and brain regions that are present in the left and right hemisphere, we compared laterality and found that, in general, the between-subject variability was much wider than the within-subject (left vs right) difference. These values were averaged to provide only a single value per structure. After splitting the data by sex, for each diffusion metric and tract, a LOESS model (span=0.75) was fit: *value ∼ age*. Then Inter Quartile Residuals (*IQR*) were computed, and visits with observed *IQR* exceeding median *3·IQR* were flagged for removal. Any subject passing these thresholds for any tract was excluded entirely. This procedure minimized technical anomalies while respecting biological heterogeneity.

### Statistical modeling of age effects

Tract/region-wise age effects on diffusion metrics (FA and MD or one of the 3 NODDI models) were modeled using Robust Linear Regression with default settings in R, MASS package, with the following model formula (*DTI|NODDI metric ∼ Age + sex*). Robust linear regression fits a linear model by using an M estimator, which uses the Huber proposal psi function to downweight the influence of outliers in iteratively reweighted least squares (IRLS). A major characteristic of RLM is that it does not make an assumption for normally distributed data. Therefore, one cannot fairly estimate a p-value. Instead, we estimate and report confidence intervals by bootstrapping the data in 1000 iterations. This approach protected inference against non-Gaussian residuals and rare measurement deviations. For each tract, we extracted β*_age_* and 95% confidence intervals (CI). **Figure S1** and **Figure 3** report the beta values (slopes of the linear model fit) obtained by fitting the RLM and the CI derived from 1000 bootstrap iterations. From this, one can fairly conclude that the effect of Age on the DTI or NODDI metric is significant if the beta value is non-null and the CI does not overlap with zero, or reject the effect of age if the CI includes zero.

### Model Selection for Linear vs Nonlinear Age Effects

For FA (**Figure 2**) and MD (**Figure S2**), linear and quadratic age models were compared using nested ANOVA-tests and AIC-based model evidence. The null model was *lm(DTI metric ∼ sex )*, the test for linear relation to age was *lm(DTI metric ∼ age + sex),* and the nonlinear fit including quadratic parameter was *lm(DTI metric ∼ poly (age, 2) + sex)*. Then we performed a nested ANOVA analysis test for linearity *anova (null, linear)* with results of the ANOVA reported in the ‘P-val linear’ column of each table where our hypothesis H_0_:“there is no significant difference between the two models” is rejected if *P-val* is less than 0.05. A second ANOVA test estimated the fit improvement by the addition of the quadratic term against the linear-only model *anova (linear, polynomial)*. The results of this second test are reported in the column ‘P-val nonlinear’ again with evidence of better fit for the polynomial being given by the estimated p-value < 0.05. One of the tracts measured for FA “Posterior corona radiata” passed the significance threshold for non-linear but did not pass the significance threshold for linear effect of age, visual inspection confirmed that there is no sufficient evidence for age effect and the non-linear conclusion is driven by outliers at older ages therefore the model choice for this tract was selected as ‘null’. Two additional metrics of evidence for non-trivial curvature were used to estimate the optimal model fit. In the first measure, we evaluated the Akaike Information Criterion (AIC) and calculated the delta AIC between the linear and the non-linear fit and set the threshold for evidence of a non-linear relation to be ΔAIC ≤ –2. For the second metric, we convert the AIC measures of the two models to relative likelihood 0 < 𝐿*_i_* ≤ 1; given model *i* = 1, K the relative likelihood of each model is defined as 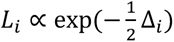 with Δ*_i_* = 𝐴𝐼𝐶*_i_* − min(𝐴𝐼𝐶) the Akaike weight or model likelihood is 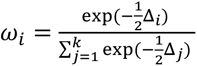 The decision rule for retaining the quadratic (nonlinear) model: if *P-val* nonlinear is *p* < 0.05 and ΔAIC ≤ −2 and/or AIC weight ≥ 0.70. If the above conditions were not satisfied but the ‘P-val linear’, of the linear age effect was *p* < 0.05 we retained the linear model. Otherwise, the model choice was classified as ‘null’ i.e. “no detectable age effect.”. This procedure allowed nonlinear aging trajectories to be detected when statistically justified.

### Canonical Correlation Analysis (CCA)

Multivariate coupling between white matter microstructure and cognitive function was evaluated using canonical correlation analysis (CCA) with the yacca package. The imaging matrix (X) comprised all 34 white matter parcels across three NODDI metrics (ficv, fiso, and odi), resulting in 102 imaging variables. The cognitive matrix (Y) comprised seven variables: spatial reaction time, verbal reaction time, memory z-score, executive control z-score, MMSE total score, animal naming, and geriatric depression scale. Both matrices were standardized to zero mean and unit variance prior to CCA, and canonical variates were computed in standardized form. To interpret the contribution of original variables to each canonical variate, structure correlations (also called canonical loadings) were computed as the Pearson correlation between each original variable and its corresponding canonical variate: for imaging variables, cor(X, U); for cognitive variables, cor(Y, V). Structure correlations represent the proportion of variance an original variable shares with the canonical variate and provides more stable interpretation than canonical coefficients, particularly when predictor variables are intercorrelated. The statistical significance of each canonical correlation was assessed using Wilks’ lambda approximation with F-tests as implemented in yacca::F.test.cca. The first canonical variate pair (CV1) showed significant correlation (R = 0.72, p < 2.2×10⁻¹⁶), explaining 52% of the shared variance between imaging and cognitive domains (**Figure S3A**). Subsequent canonical variates (CV2-CV7) did not reach statistical significance after correction for multiple comparisons and are reported for completeness but not interpreted (**Figure S3C**). To assess whether age contributed to the structure-cognition relationships captured by CCA, each imaging canonical variate (U1-U7) and cognitive canonical variate (V1-V7) was regressed against age using ordinary least squares regression. P-values were adjusted for multiple comparisons across all 14 variates using the false discovery rate (FDR) method. Standardized regression coefficients (β), 95% confidence intervals, and FDR-adjusted p-values are reported (**Figure S3B and S3C**). To visualize the structure in the data, pairwise Pearson correlations were computed for between the X and Y matrices (**Figure S3E**) within the imaging matrix X (**Figure S3G**) and the cognitive matrix Y (**Figure S3F**).

## Acknowledgements

We acknowledge the support of NIH NIA P01AG066591 and T32-AG000266 (PI: Lisa Ellerby). D.F. and L.M.E. acknowledge funding from Hevolution Foundation.

## Authors Contributions

Conceptualization: NTM, IMG, Methodology and Data Collection: NTM, JK, KC, MO, Investigation: NTM, IMG, LME, Visualization: NTM, IMG, LME, Funding acquisition: LME, Project administration: LME, DF, Supervision: LME, DF, Writing-original draft: NTM. Writing-review and editing: NTM, LME, IMG, JK, KC

## Competing Interests

## Supplementary figure legends

**Supplementary Figure 1.**
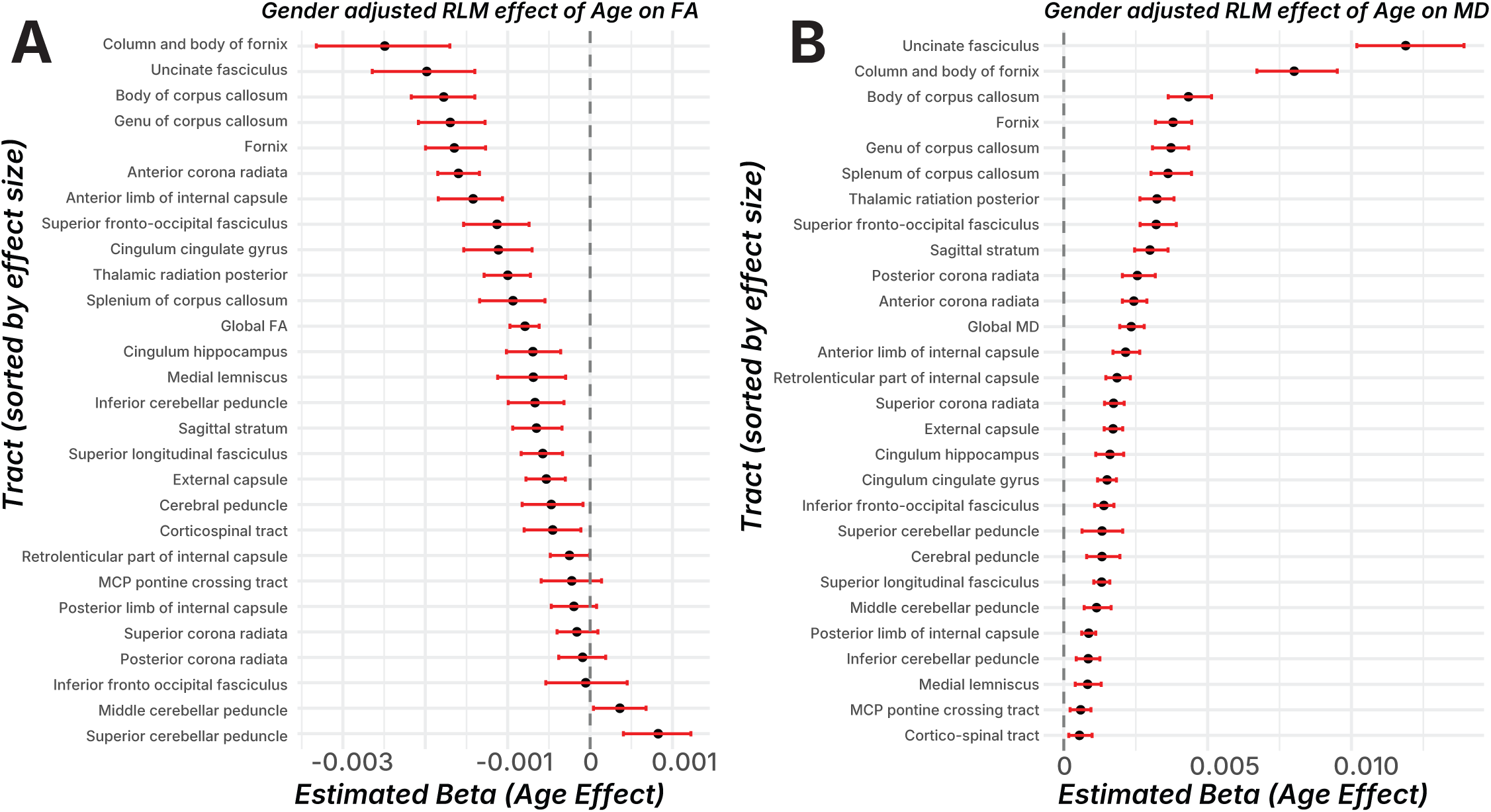
Participant age is a major factor influencing dwMRI measures (FA and MD). **(A)** Magnitude of the beta estimates of robust linear model (RLM), fitting the effect of age on fractional anisotropy (FA) for each white-matter tract, adjusted for sex (FA ∼ Age + sex). For every tract, FA values were derived from the first available dwMRI visit, left–right hemisphere homologs were averaged, and scans with robust LOESS-based outlier patterns were removed at the participant level. Age effects were estimated using a resistant linear model and statistical uncertainty was quantified by non-parametric bootstrapping (1,000 resamples). Points represent the estimated age coefficient (β), and horizontal red bars denote the bootstrapped 95% percentile confidence interval. Tracts are ordered by decreasing magnitude of age effect. The dashed vertical line indicates β=0 (i.e. no effect of age). Negative values indicate age-related decreases in FA. **(B)** Magnitude of the beta estimates of robust linear model (RLM), fitting the effect of age on mean diffusivity (MD) for each white-matter tract, adjusted for sex (MD ∼ Age + sex). For every tract, MD values were derived from the first available dwMRI visit, left–right hemisphere homologs were averaged, and scans with robust LOESS-based outlier patterns were removed at the participant level. Age effects were estimated using a resistant linear model and statistical uncertainty was quantified by non-parametric bootstrapping (1,000 resamples). Points represent the estimated age coefficient (β), and horizontal red bars denote the bootstrapped 95% percentile confidence interval. Tracts are ordered by decreasing magnitude of age effect. The dashed vertical line indicates β=0 (i.e. no effect of age). Positive values indicate age-related increases in MD.

**Supplementary Figure 2.**
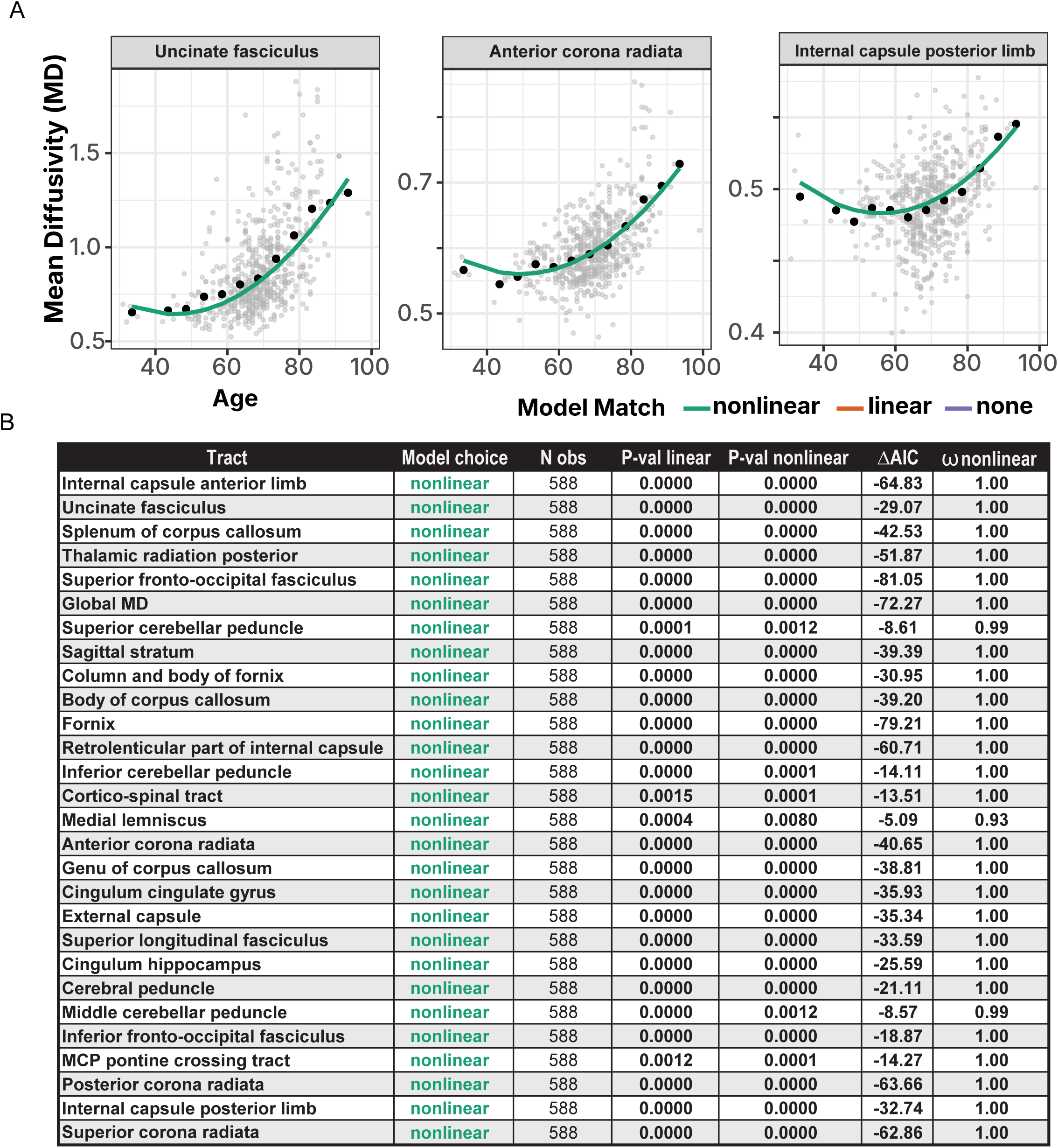
Age-related trajectories of Mean diffusivity (MD) across major white matter pathways show heterogenous accelerations. **(A)** Representative scatter plots of mean tract values from first visit dwMRI scans (gray points) MD against chronological age. Large black points indicate the mean MD within a 5-year interval. For each tract, age trajectories were estimated using sex adjusted regression models; the best-fitting model was selected using a combined statistical and information-theoretical criterion (see table in panel B). Colored curves denote the chosen model for each tract: green = nonlinear (quadratic) age dependence; orange = linear age dependence; purple = non-detectable age effect. Panels are scaled independently to emphasize tract-specific dynamic ranges. **(B)** Table of model fitting diagnostics for each pathway. **Tract**: white matter pathway of interest. **Model choice**: best supported age model (nonlinear (green), linear (orange), none (purple), based on significance testing and information criterion evidence. **N obs:** number of individuals contributing MD measurements to that tract. **P-val linear**: significance score of nested ANOVA F test of the linear age effect (*MD∼Age + sex*) versus a gender-only null model (*MD∼sex*). **P-val nonlinear**: significance of nested ANOVA F test for curvature (quadratic term) (*MD ∼poly(Age, 2) + sex*) beyond the linear model (*MD∼Age + sex*); **ΔAIC**: Akaike Information Criterion (AIC) difference between linear and nonlinear term, if (ΔAIC ≤ –2, and/or ω ≥ 0.70) are labeled nonlinear; those with significant but non-curved age effects are labeled linear; and tracts without significant age dependence are labeled none. **ω nonlinear** Akaike weight for the nonlinear model, representing the relative probability that curvature provides the best description of age-related change.

**Supplementary Figure 3.**
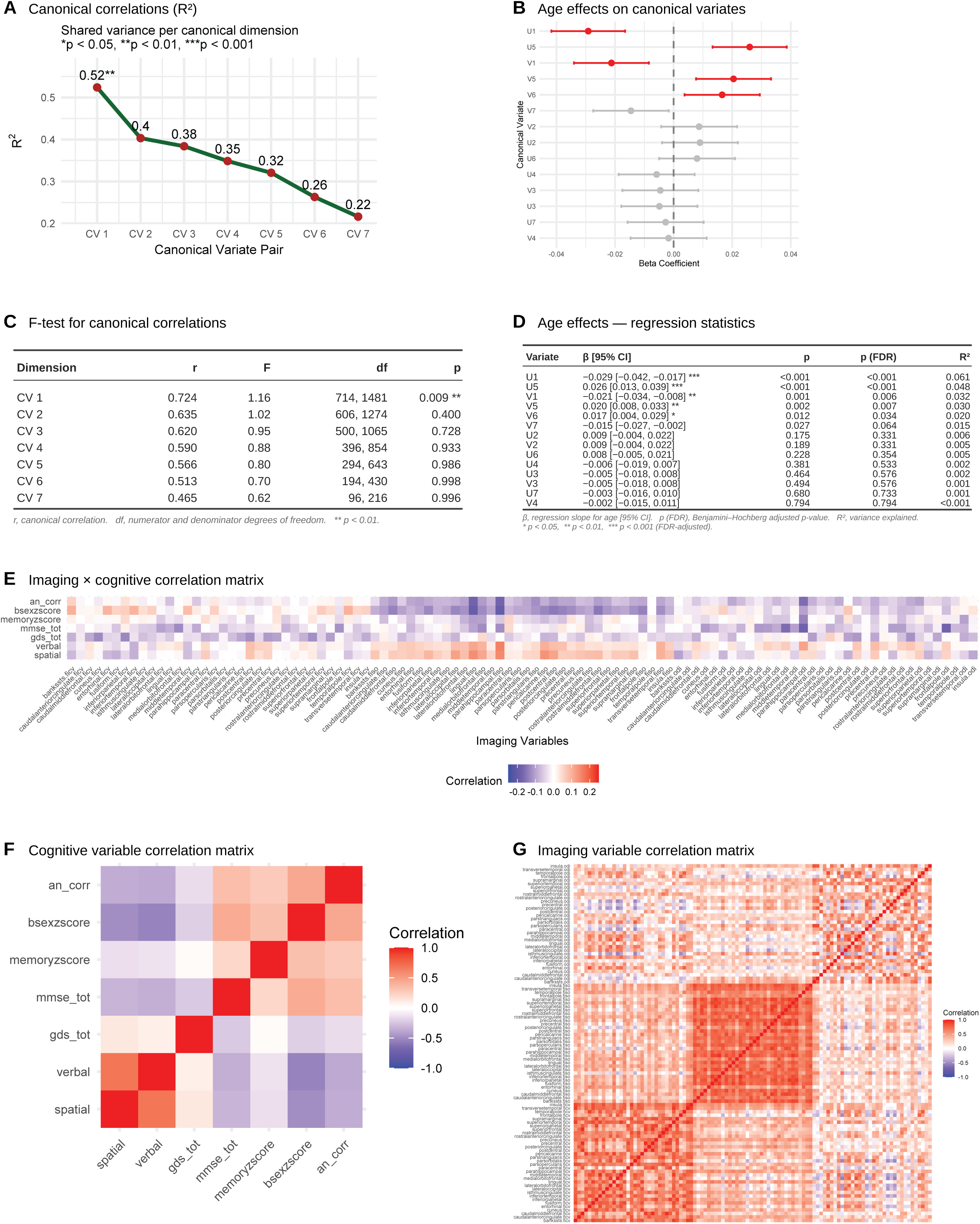
Multivariate associations between white matter microstructure and cognition show age significantly impacts the dominant CCA dimension. Canonical correlation analysis (CCA) was applied to multi-tract neurite density indices and composite cognitive scores to identify latent modes of shared variance. Wilks’ lambda–based F-test showed statistical significance for the dominant dimension but not the remaining ones. **(A)** Scree plot of the 7 canonical variates, shared variance (R2) between each pair of canonical variates. **(B)** Regression of each canonical variate on age (linear models with FDR correction), shown with regression coefficients and 95% confidence intervals. For the significant canonical variate CV1 both the imaging variate (U) and the cognitive variate (V) are significant (red) while individually U5, V5 and V6 also associated significantly with age they are not part of a significant canonical variate (CV). **(C)** Table of statistical significance tests using the Rao’s approximation of F show CV1 to be the only statistically significant dimension. **(D)** Statistical summary table of the regression of all the canonical variates against age (presented in panel B). **(E)** The correlation matrix between the two datasets. The NODDI measures are grouped by model (ficv, fiso, odi). Note the visibly high correlation values with cognition across the fiso model. **(F)** The correlation matrix of cognitive variables. **(G)** The correlation matrix of the NODDI measures shows little structure within each model.

## Notes

### Competing Interest Statement

The authors have declared no competing interest.

